# Luminal Flow in the Connecting Tubule induces Afferent Arteriole Vasodilation

**DOI:** 10.1101/2024.09.12.612758

**Authors:** Hong Wang, Pablo A. Ortiz, Cesar A. Romero

**Author notes:** Corresponding author: Cesar A. Romero, M.D., Renal Medicine Division, Department of Medicine, Emory University School of Medicine, 101 Woodruff Circle, Woodruff Memorial Research Building, Office 338A, Atlanta, Georgia, 30322., Phone-FAX : (404) 727-395.

## Abstract

**Background:** Renal autoregulatory mechanisms modulate renal blood flow. Connecting tubule glomerular feedback (CNTGF) is a vasodilator mechanism in the connecting tubule (CNT), triggered paracrinally when high sodium levels are detected via the epithelial sodium channel (ENaC). The primary activation factor of CNTGF—whether NaCl concentration, independent luminal flow, or the combined total sodium delivery—is still unclear. We hypothesized that increasing luminal flow in the CNT induces CNTGF via O2^-^ generation and ENaC activation.

**Methods:** Rabbit afferent arterioles (Af-Arts) with adjacent CNTs were microperfused *ex-vivo* with variable flow rates and sodium concentrations ranging from <1 mM to 80 mM and from 5 to 40 nL/min flow rates.

**Results:** Perfusion of the CNT with 5 mM NaCl and increasing flow rates from 5 to 10, 20, and 40 nL/min caused a flow rate-dependent dilation of the Af-Art (p<0.001). Adding the ENaC blocker benzamil inhibited flow-induced Af-Art dilation, indicating a CNTGF response. In contrast, perfusion of the CNT with <1 mM NaCl did not result in flow-induced CNTGF vasodilation (p>0.05). Multiple linear regression modeling (R^2^=0.51;p<0.001) demonstrated that tubular flow (β=0.163 ± 0.04;p<0.001) and sodium concentration (β=0.14 ± 0.03;p<0.001) are independent variables that induce afferent arteriole vasodilation. Tempol reduced flow-induced CNTGF, and L-NAME did not influence this effect.

**Conclusion:** Increased luminal flow in the CNT induces CNTGF activation via ENaC, partially due to flow-stimulated O2-production and independent of nitric oxide synthase (NOS) activity.

## Introduction

Glomerular filtration rate (GFR) is maintained constant by three renal autoregulatory mechanisms that modulate renal resistance and glomerular pressure: the myogenic response, tubuloglomerular feedback in the macula densa (TGF), and connecting tubule glomerular feedback (CNTGF) in the connecting tubule (CNT)[1,2]. CNTGF is a vasodilator mechanism initiated in the CNT via epithelial Na+ channels (ENaC)[3–5]. ENaC senses Na+ in the CNT and triggers the release of prostaglandins and epoxyeicosatrienoic acids from the principal cells of the CNT, which paracrinally induce afferent arteriole vasodilation, leading to increased renal blood flow and GFR[2]. Thus, a high amount of sodium present distally in the CNT indicates the need to increase the sodium-filtered load by inducing renal vasodilation and promoting Na+ excretion, which is the central role of CNTGF[5].

Luminal flow in the nephron varies due to GFR and fluid reabsorption[6,7]. In rabbits, single nephron GFR has been reported to be close to 20 nL/min, with more than half reabsorbed in the proximal tubule, indicating that less than 10 nL/min can be observed in the distal nephron under physiological conditions[8]. However, luminal flow can increase significantly under some pharmacological or pathological circumstances [9,10]. Sodium concentration in the CNT is highly variable according to body needs, intake, and reabsorption in proximal segments, usually ranging from <10 mM to 80 mM and rarely exceeding 100 mM[8,11]. Previous literature, including our studies, has explored the activation of CNTGF with sodium concentrations ranging from 10 mM to 80 mM while keeping the perfusion rate constant at 15 nL/min[3,11,12].

Morimoto et al. speculated that hydrodynamic forces associated with increased urinary flow rate activate ENaC[13]. Satlin and Kleyman have shown that luminal flow modifies ENaC activity [14]. Thus, there is evidence that flow can activate ENaC. Consequently, we speculate that flow can activate CNTGF. In this work, we hypothesize that increasing luminal flow in the CNT induces CNTGF via ENaC activation.

Superoxide (O2^-^) and nitric oxide (NO) are physiological regulators of renal function[15,16]. Increases in luminal flow stimulate O2^-^ production[17,18], and O2^-^ activates ENaC in the cortical collecting duct (CCD)[19,20]. Conversely, NO is an ENaC inhibitor and is also stimulated by luminal flow in the kidney [21]. The roles of O2^-^ and NO in the CNT and flow-mediated CNTGF are unknown. We propose that luminal flow-stimulated O2^-^ acts in an autocrine manner in the CNT to activate Na+ transport by ENaC, thus participating in flow-induced CNTGF.

To test these hypotheses, we used an *ex-vivo* technique involving simultaneous perfusion of a microdissected rabbit Af-Art and adherent CNT, thereby avoiding the confounding influence of multiple systemic factors that regulate renal microcirculation. Using this preparation, we found that luminal flow has an independent effect on inducing dilation of Af-Art. This effect depends on ENaC activation, which requires a minimum amount of sodium. We also found that the flow-induced CNTGF response is partly due to flow-stimulated O2^-^ production by the CNT.

## Materials and methods

All experiments were approved by the Henry Ford Health System Institutional Animal Care and Use Committee (IACUC) and were conducted in accordance with the National Institutes of Health Guidelines for the Care and Use of Laboratory Animals. We used methods similar to those previously described to isolate and microperfuse the afferent arteriole (Af-Art) with attached glomerulus and connecting tubule (CNT)[3,22]. Male New Zealand white rabbits (Covance Research Products, Kalamazoo, MI, USA; 1.5 to 2.5 kg) were fed standard rabbit chow with 0.34% Na+ and 0.40% Cl-(Ralston Purina, St. Louis, MO, USA) and given tap water ad libitum. Rabbits were anesthetized with ketamine plus xylazine (50 mg/kg and 10 mg/kg IM), sodium pentobarbital (30 mg/kg IV). Heparin (500 U IV) was injected to prevent blood coagulation. The kidneys were removed and sliced along the longitudinal corticomedullary axis. Slices were placed in ice-cold minimum essential medium (Gibco Laboratories, Grand Island, NY, USA) containing 5% bovine serum albumin (Sigma, St. Louis, MO, USA) and dissected under a stereomicroscope (SZH; Olympus) (Figure 1A). A superficial Af-Art with the adherent CNT was dissected from each rabbit using fine forceps as previously described[3]. The microdissected complex was transferred to a temperature-regulated perfusion chamber mounted on an inverted microscope using a micropipette. Both the Af-Art and CNT were cannulated with an array of glass pipettes (Figure 1B), as described previously[3]. The Af-Art was perfused with a minimum essential medium containing 5% bovine serum albumin. The medium was oxygenated with room air. Intraluminal pressure was measured by Landis’ technique and maintained at 60 mmHg[23]. The CNT was perfused using a nanoliter infusion pump with variable NaCl concentrations and flow rates in different protocols (see below). The primary solution contained (in mM) 10 HEPES, 1 CaCO3, 0.5 K2HPO4, 4 KHCO3, 1.2 MgSO4, 5.5 glucose, 0.5 sodium acetate, and 0.5 sodium lactate (pH 7.4). The perfusion rate of the CNT was from 5 to 10, 20, and 40 nL/min. The bath was superfused with a minimum essential medium containing 0.15% bovine serum albumin at a 1 ml/min rate. Microdissection and cannulation of the Af-Art and tubular segment were completed within 90 min at 8°C, after which the temperature was gradually raised to 37°C. A 30-minute equilibration period was allowed before taking any measurements. Images of the Af-Art were displayed at magnifications up to 1980X. As isolated arterioles have little or no tone, the studies were performed in 0.2-0.5 µM norepinephrine (NE). Preconstricted Af-Arts decreased basal diameter by 46-48% of resting diameter. The Af-Art diameter was measured in the region of maximal response to NE at three sites separated by 3–5 μm and expressed as the average of these three measurements (Figure 1B). As we previously reported, the diameter was recorded at 5-sec intervals with a video camera and measured using a computer equipped with a Metavue image analysis system[11]. All protocols consisted of two consecutive dose-response curves in the same preparation, where the Af-Art diameter was measured at each point. One dose-response curve served as a time control and the other as an experimental. The first (Flow) protocol included a dose-response curve to a fixed 5mM NaCl concentration with increasing flow rates (5, 10, 20, and 40 nL/min). The second protocol (<u>No-Salt protocol</u>) was a dose-response curve with less than 1mM NaCl to evaluate the effect of increased flow rate without NaCl. The third (<u>Concentration protocol</u>) consisted of a 10 nL/min fixed flow with two NaCl concentrations (10mM and 80mM). Finally, the fourth protocol evaluated the role of O2- and NO on flow-mediated vasodilation by infusing Tempol (100 μM) in the CNT with and without L-NAME (100 μM) while increasing flow from 5 to 40 nL/min with a fixed 5mM NaCl concentration. Combining all the data, we obtained measurements with a broad spectrum of physiological and non-physiological conditions, including low, medium, and high luminal flow (5 to 40 nL/min), with low, medium, and high NaCl concentrations (<1mM, 5mM, 10mM, and 80mM). Sodium delivery (NaV) was calculated by multiplying sodium concentration (mM) times luminal flow (nL/min) and expressed as nmol/min. Combined data from all protocols were used to build a table with 70 individual observations of flow, NaCl concentrations, and the corresponding NaV to model afferent arteriole vasodilation. L-NAME, benzamil, and Tempol (4-Hydroxy-TEMPO) were purchased from Sigma.

**Figure 1.**
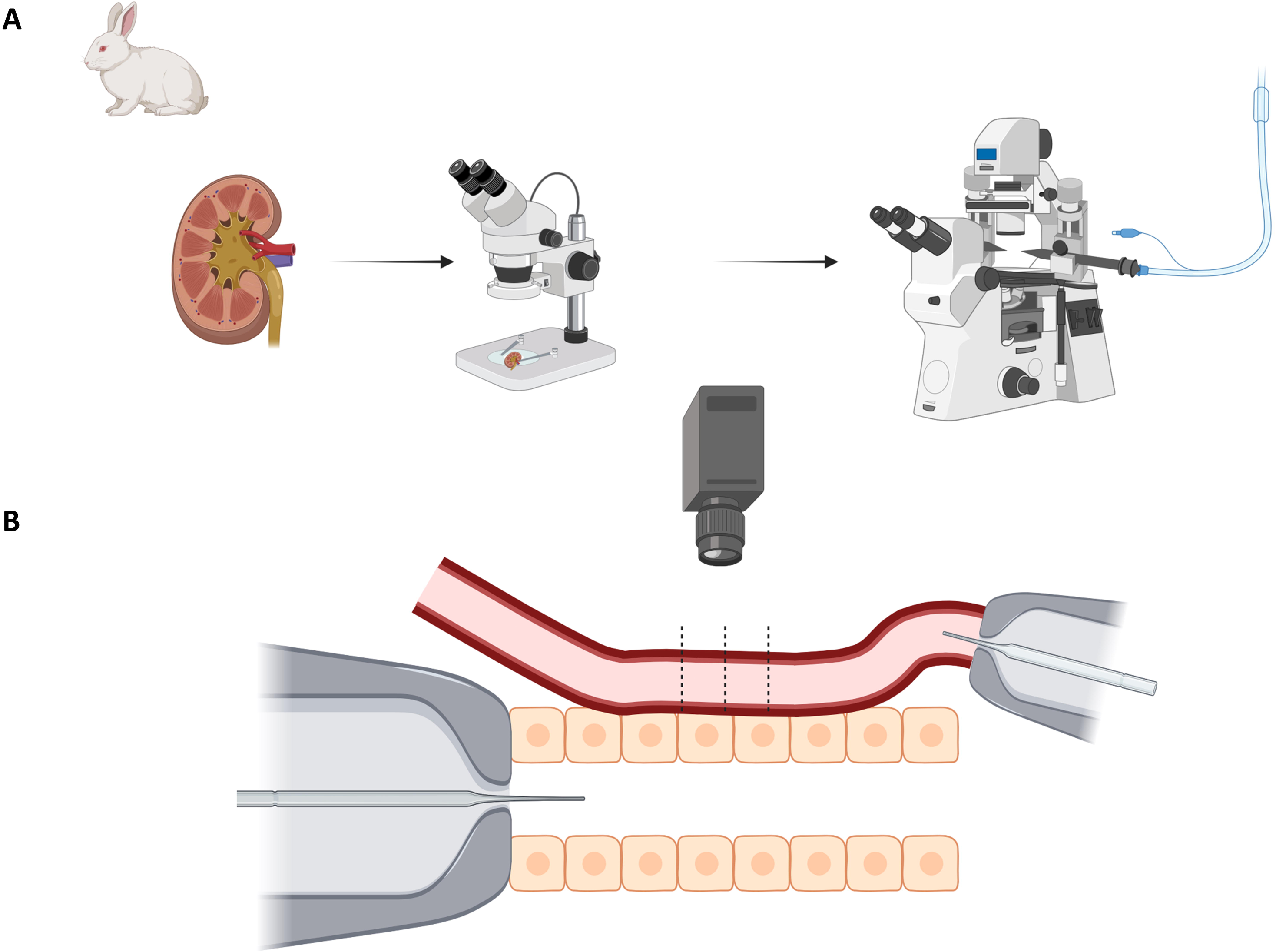
Kidney tubule microperfusion. Male New Zealand white rabbits were anesthetized, and their kidneys were removed and sectioned along the corticomedullary axis. Superficial afferent arterioles (Af-Art) with attached connecting tubules (CNT) were microdissected under a stereomicroscope. The isolated complexes were transferred to a temperature-controlled perfusion chamber and viewed using an inverted microscope. Both the Af-Art and CNT were cannulated and perfused with glass pipettes. The CNT was perfused with variable NaCl concentrations and flow rates. Af-Art diameter was measured at three points, averaged, and recorded at 5-second intervals. Each protocol involved two consecutive dose-response curves. Measurements were analyzed using Metavue software.

### Statistical considerations

Data are expressed as mean ± SE. When comparing two dose-response curves, we determined whether the curves differed and whether the individual points in one curve differed from their equivalents in the other. The students’ paired t-test was used to compare the paired differences for each two-group comparison. When multiple comparisons were performed, Hochberg’s step-up procedure was used to adjust the P values. The type I error rate was set at 0.05.

A multiple linear regression model was built to determine the effect and weight of luminal flow, NaCl concentration, and sodium delivery (NaV) on Af-Art diameter. In the model, Af-Art diameter was the dependent variable, and luminal flow, NaCl concentration, and NaV were the independent variables. Multicollinearity was tested through correlation and variance inflation factors. The model fit was evaluated through the R^2^ coefficient and mean square error (MSE). Several models were run using stepwise criteria, and the full model was chosen as the best–performing one.

## Results

We first performed a dose-response curve to measure the flow rate while measuring the Af-Art diameter. The resting Af-Art diameter, when the CNT was perfused with a low perfusion rate (5 nL/min) and 5 mM NaCl concentration, was 16.7 ± 0.3 µm. This decreased to 9.8 ± 0.5 µm by adding NE to the perfusion bath. Pre-constriction with NE is necessary to study vasodilation and confirm the isolated arteriole’s vitality. The Af-Art then dilated to 10.4 ± 0.6, 12.5 ± 0.4, and 14.8 ± 0.4 µm when the flow rate in the CNT was increased to 10, 20, and 40 nL/min, respectively, showing that with a constant NaCl concentration, increasing luminal flow in the CNT dilates the Af-Art (Figure 2A). We then returned the CNT flow rate to 5 nL/min and performed a second dose-response curve to flow. Pre-constricted Af-Art diameter was initially 9.7 ± 0.5 µm when the CNT was perfused at 5 nL/min, then it dilated to 10.6 ± 0.5, 12.9 ± 0.3, and 14.8 ± 0.3 µm at 10, 20, and 40 nL/min, respectively (n = 7; Figure 2A). Thus, the flow response to increased luminal flow was reversible, reproducible, and time-independent.

**Figure 2.**
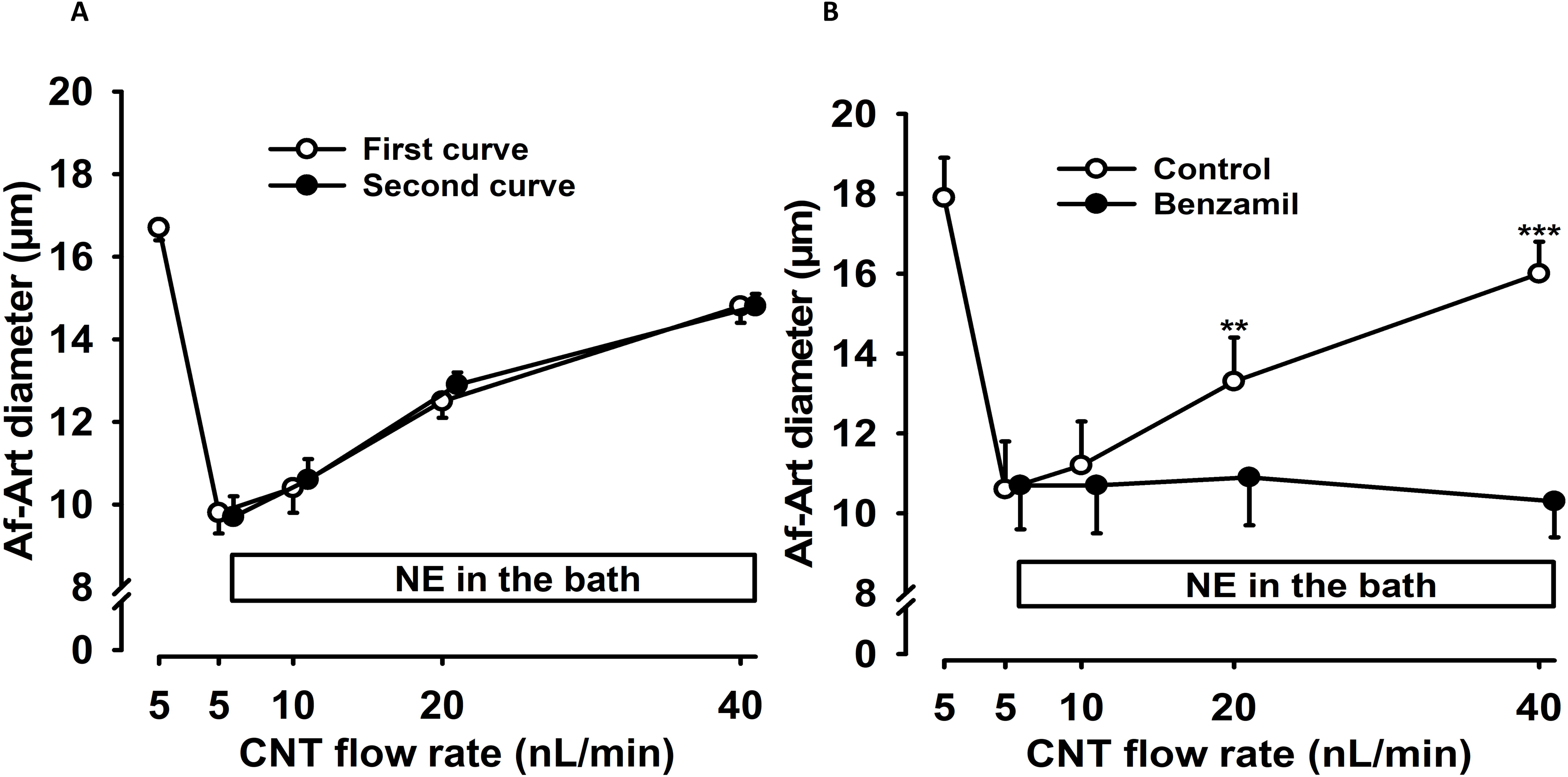
Luminal flow in the connecting tubule induces afferent arteriole dilation. The connecting tubule (CNT) was perfused with 5 mM NaCl. Luminal flow rates of 10, 20, and 40 nL/min induced significant vasodilation of preconstricted afferent arteriole (Af-Art). Figure 2A shows no difference between the first (O) and second (•) dose-response curves (n = 7), indicating the reproducibility of vasodilation and independence from time (time control). Figure 2B demonstrates the effect of benzamil in the CNT on flow-induced Af-Art dilation. Increasing the CNT flow rate significantly dilated the Af-Art (O), but 0.1 µM benzamil (•) in the CNT completely inhibited this effect (n = 6), indicating the participation of the ENaC-dependent connecting tubule glomerular feedback. ***p < 0.001, benzamil vs. control. NE = norepinephrine; CNT = connecting tubule; Af-Art= Afferent arteriole.

To determine whether ENaC activation is involved in flow-induced Af-Art dilation, the ENaC inhibitor benzamil (0.1 µM) was added to the CNT perfusate to block ENaC (Figure 2B). In the control curve, Af-Art diameter increased from 10.6 ± 1.2 to 11.2 ± 1.1, 13.3 ± 1.1, and 16.0 ± 0.8 µm when CNT luminal flow was raised from 5 to 10, 20, and 40 nL/min, respectively. In the presence of benzamil, Af-Art diameter did not change (n= 6; p < 0.001, control vs. benzamil; Figure 2B). These results confirm that Af-Art dilation induced by changes in CNT luminal flow is due to CNTGF.

To determine whether NaCl was necessary for the CNTGF response to flow, we first perfused the CNT with less than 1 mM NaCl concentration. We found that increasing the CNT flow rate did not induce changes in Af-Art diameter (Figure 3A). When the CNT perfusate was elevated to 5 mM NaCl, increases in the CNT flow rate from 5 to 10, 20, and 40 nL/min caused significant dilation of the Af-Arts (Figure 3A; n = 6; p < 0.001, <1 mM NaCl vs. 5 mM NaCl). These data indicate that a minimum NaCl concentration is necessary for a CNTGF response to flow, activating ENaC in the CNT.

**Figure 3.**
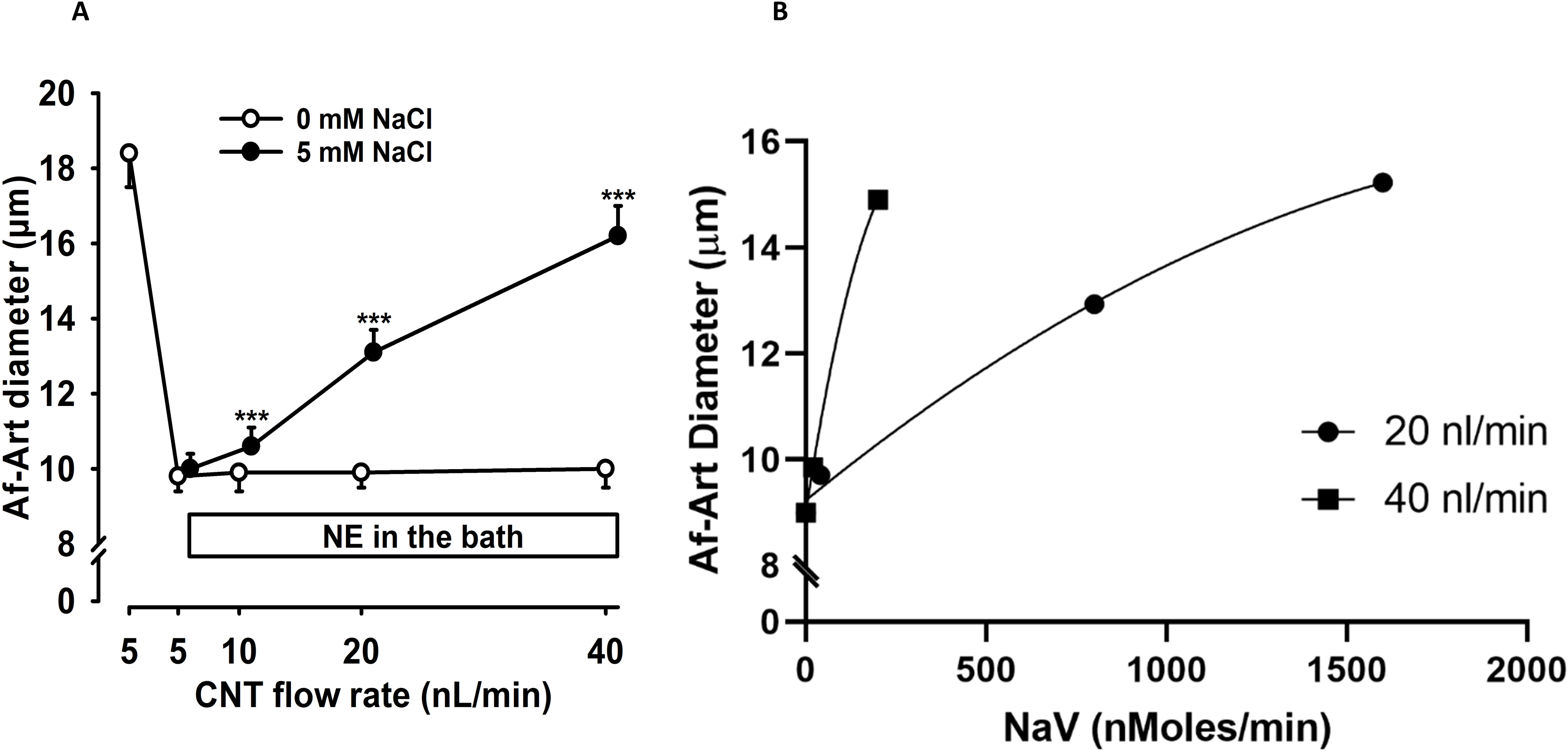
Effect of luminal flow on NaCl-induced vasodilation. When the CNT was perfused with 5 mM NaCl, increasing the flow rate significantly dilated the afferent arteriole (Af-Art) (•). However, no dilation occurred when the CNT was perfused with 0 mM NaCl (O) (n = 6), indicating the dependence of sodium for the flow-induced vasodilation (Figure 3A). ***p < 0.001, 0 mM NaCl vs. 5 mM NaCl. Figure 3B plots Af-Art diameter against sodium delivery (NaV) at two luminal flow rates. The 40 nL/min slope was steeper than at 20 nL/min, indicating an independent effect of luminal flow, adjusted for NaV, on Af-Art diameter. NE = norepinephrine; CNT = connecting tubule; Af-Art= Afferent arteriole.

Despite keeping sodium chloride concentration constant, changes in flow induce changes in sodium delivery (NaV). To explore whether luminal flow, independent of sodium concentration and sodium delivery, affects Af-Art diameter, we modeled Af-Art diameter at different levels of luminal flow adjusted by NaV (Table 1). The model explains 57% of the Af-Art variability (R² = 0.57), and the MSE was 5.7. Table 1 shows the parameter estimates. Both luminal flow and NaCl concentration were independent determinants of Af-Art diameter adjusted by NaV (p < 0.001). Figure 3B illustrates data of two different flow rates adjusted by NaV on Af-Art diameter. The slope of the curve is steeper when the high flow (40 nL/min) is compared to the standard/low flow (20 nL/min).

**Table 1.** Afferent Arteriole Diameter Modeling to Luminal Flow and Sodium Delivery (NaV).

| Variable | Parameter estimate ($\beta$) | SE | P value* |
| --- | --- | --- | --- |
| Intercept | 6.49 | 0.85 | <0.001 |
| Flow (nl/min) | 0.163 | 0.04 | <0.001 |
| NaCl (mM) | 0.140 | 0.03 | <0.001 |
| NaV (nMoles/min) | -0.004 | 0.002 | 0.08 |

\*Multiple Linear regression model $R^2=0.57$ MSE 5.7

To study the mechanism associated with flow-induced vasodilation, we tested the role of increased luminal flow-stimulated O2^-^ on CNTGF. The O2^-^ dismutase mimetic tempol (100 µM) was added to the CNT perfusate. In the control dose-response curve to flow, Af-Art diameter increased from 9.8 ± 0.5 to 10.8 ± 0.5, 13.6 ± 0.5, and 16.2 ± 0.7 µm when the CNT flow rate was raised from 5 to 10, 20, and 40 nL/min. In the presence of tempol, the diameter increase was blunted (n = 6; p < 0.001, control vs. tempol; Figure 4A). Tempol in the CNT significantly reduced luminal flow-induced Af-Art dilation, suggesting that increased luminal flow-stimulated O2^-^ in the CNT participates in the flow-induced CNTGF response.

**Figure 4.**
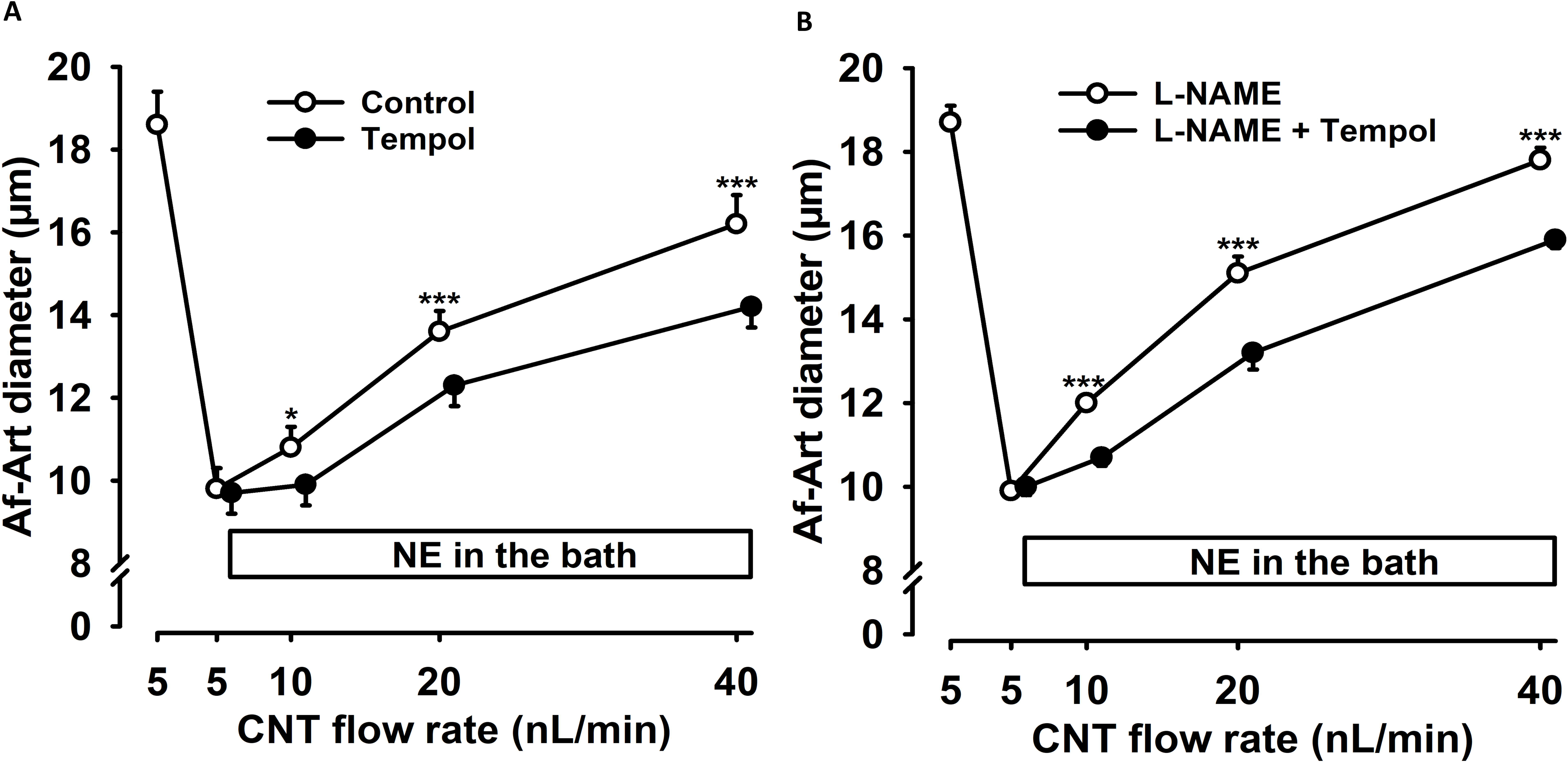
Effect of O ^-^ on flow-induced CNTGF. Increasing the CNT luminal flow rate significantly dilated the Af-Art (O), but the addition of 100 µM tempol (•) in the CNT significantly attenuated this response (Figure 4A, n = 6, *p < 0.05; ***p < 0.001, tempol vs. control). Figure 4B shows that adding 100 µM L-NAME to the tempol infusion further attenuated vasodilation (•) compared to L-NAME alone, suggesting that the effect of O2⁻ is independent of its action on NO (n = 6). ***p < 0.001, L-NAME vs. L-NAME + tempol. NE = norepinephrine; CNT = connecting tubule. NE = norepinephrine; CNT = connecting tubule; Af-Art= Afferent arteriole.

To study whether the effect of O2^-^ on flow-induced CNTGF was dependent on bioavailable nitric oxide (NO), we examined the impact of tempol on flow-induced CNTGF in the presence of an NO synthase (NOS) inhibitor, L-NAME (100 µM). In the control dose-response curve to flow, L-NAME was added to the CNT perfusate, and Af-Art diameter increased from 9.9 ± 0.2 to 12.0 ± 0.2, 15.1 ± 0.4, and 17.8 ± 0.3 µm when the flow was raised from 5 to 10, 20, and 40 nL/min. The diameter increase was blunted in the presence of L-NAME plus tempol (n = 6; p < 0.001, L-NAME vs. L-NAME + tempol; Figure 4B). These results suggest increased flow-stimulated O2^-^ production participates in flow-induced CNTGF independent of NO.

## Discussion

Using isolated perfused Af-Art and attached CNT, we have obtained evidence that luminal flow in the CNT independently affects Af-Art vasodilation through CNTGF activation. The Af-Art dilation appears to be initiated by Na+ absorption in the CNT because when we perfused the CNT with <1 mM NaCl, we could not induce a CNTGF response. We also found that blocking ENaC in the CNT with benzamil completely inhibited the dilation of the Af-Art. We performed regression modeling to identify the independent effect of luminal flow, NaCl concentration, or the combined sodium delivery.

We confirmed that changes in luminal flow are independent of sodium concentration and sodium delivery in inducing Af-Art vasodilation. The data indicate that CNTGF may be more sensitive to flow increases than NaCl concentration increases (β = 0.163 vs. β = 0.140, respectively; p < 0.001). Thus, we can achieve maximal vasodilation by doubling the luminal flow from 20 to 40 nL/min. However, the NaCl concentration must increase more than eight times to reach maximum vasodilation. The present results indicate that in flow-induced CNTGF, a small amount of Na+ is needed. The minimum amount of sodium that activates CNTGF tested in previous reports was 5 mM at ∼20 nL/min[3,4]. In the present study, we tested changes in flow as low as 5 nL/min with a 5 mM NaCl concentration. However, the minimum NaCl needed to induce CNTGF activation must be determined in future studies and can be between 1 and 5 mM, according to the present study.

Previous studies suggest that increases in tubular flow rate activate ENaC, stimulating Na+ absorption in the cortical collecting duct[14,24]. Satlin et al.[14] examined the effect of increases in fluid flow rate on exogenous ENaC channels expressed in X. *laevis* oocytes and endogenous amiloride-sensitive Na+ channels in rabbit CCDs. Their results suggest that ENaC is a flow-regulated ion channel. Increased luminal flow can activate ENaC by exerting three distinct mechanical forces on epithelial cells in a tubule: 1) shear stress, 2) stretch, and 3) pressure. It may also be activated by enhanced ion delivery. Our current results indicate that flow in the CNT activates ENaC, which is prevented with benzamil, inducing the CNTGF response. This aligns with previous studies where luminal flow in the distal nephron increases ENaC activity[13]. We further found that the O2^-^ dismutase mimetic tempol in the CNT significantly reduced CNTGF and that inhibition of NOS with L-NAME did not block the effect of tempol, suggesting that the impact of O2^-^ was not related to a NO scavenging effect. O2^-^ is a physiological regulator of renal function, promoting salt and water retention, and has been implicated in salt-sensitive hypertension [25,26]. Increases in luminal flow stimulate O2^-^ production [18,27], and O2^-^ activates ENaC in the CCD[19]. We previously showed that O2^-^ mediates the enhancement of CNTGF by luminal ANG II acting downstream from protein kinase[12]. However, the mechanisms of O2^-^ activating CNTGF are still unknown and need further investigation.

Flow-induced CNTGF activation may be relevant in conditions with increased distal sodium delivery, such as glomerular hyperfiltration in diabetes or the remnant kidney after unilateral nephrectomy, or with medications such as diuretics. Flow-induced CNTGF activation may negatively affect low-flow situations such as cardiorenal syndrome, dehydration, or hepatorenal syndrome. Future research must address these situations and their link to renal hemodynamic regulation.

In conclusion, our studies demonstrate the independent effect of luminal flow on Af-Art vasodilation by activating CNTGF. Flow-induced O2-produced by the CNT positively regulates CNTGF by stimulating Na+ transport.

## Authors Contributions

H. Wang and CA Romero conceptualized the study and were responsible for data acquisition, analysis, and interpretation. P.A. Ortiz was responsible for the analysis and interpretation of the data obtained. H. Wang and C.A. Romero wrote the original manuscript. C. A Romero and P.A. Ortiz were responsible for funding acquisition, methodology, project administration, resources, and supervision. All authors reviewed and edited the manuscript.

## Compliance with Ethical Standards

The authors have no financial conflict of interest to disclose. All experiments were conducted in accordance with the National Institutes of Health Guidelines for the Care and Use of Laboratory Animals and approved by the Institutional Animal Care and Use Committee (IACUC).

## Data Sharing Statement

All data used in this study are available at the authors’ reasonable request.

